# Simulating human-in-the-loop optimization of exoskeleton assistance to compare optimization algorithm performance

**DOI:** 10.1101/2024.04.05.587982

**Authors:** Zoe Kutulakos, Patrick Slade

**Affiliations:** School of Engineering and Applied Sciences, Harvard University, Cambridge MA, USA

## Abstract

Assistive robotic devices like exoskeletons offer the promise of improving mobility for millions of people. However, developing devices that improve an objective mobility metric is challenging. Human-in-the-loop optimization is a systematic approach for personalizing robotic assistance to maximize a mobility metric that has improved device performance for different metrics and applications. Successfully performing human-in-the-loop optimization requires the experimenter to make many decisions, like selecting the appropriate optimization algorithm, hyperparameters, and convergence criteria. Typically, selecting these experimental settings involves pilot experimentation. We propose an approach that uses a probabilistic surrogate model, mapping assistance parameters to corresponding experimental evaluations of the objective mobility metric, to simulate human-in-the-loop optimization and inform these decisions. In this paper, we form a surrogate model of the metabolic landscape of walking with exoskeleton assistance using an existing experimental dataset. We simulate human-in-the-loop optimization by using a synthetic metabolic landscape model to evaluate the metabolic cost of walking with different assistance parameters, instead of performing an experimental measurement. We perform three simulated scenarios optimizing assistance for an expert subject, a novice subject adapting to the device, and an expert subject with up to 20 assistance parameters. The code and analyses from this work are open-source to promote use by other researchers. Simulation enables direct comparison of optimization settings to inform experimental human-in-the-loop optimization and potentially reduce the resources and time required to develop effective assistive devices.

## I. Introduction

Mobility challenges impact the lives of 1 in 7 adults, with major repercussions on physical health, mental health, independence, and quality of life [1]. Assistive robotic devices like exoskeletons offer a potential solution to mobility challenges but most exoskeletons do not yet provide benefits in real-world settings [2], [3]. Iterative exoskeleton controller development has led to steady improvements in device performance evaluated in laboratory settings [4]. Nevertheless, developing effective devices remains challenging due to the complexities of human-robot interaction. Biomechanics simulation captures important information about the musculoskeletal system and dynamics of a person [5], but simulations of the human’s biomechanical response to assistance differs from experimental results [6].

Human-in-the-loop optimization is an approach for systematically tuning robotic assistance, which can improve exoskeleton performance [7]. During optimization, a subject uses a robotic device, like an exoskeleton, and the device’s performance metric, such as metabolic cost, is directly measured. An optimization algorithm uses these performance measurements, corresponding to different assistance parameters, to inform the selection of the next parameters to experimentally evaluate. This optimization continues until reaching convergence criterion of not detecting further improvements in the performance metric within a certain period of time, or evaluating a maximum number of parameter sets. Human-in-the-loop optimization does not use an explicit model of the human or robotic system, relying instead on the experimental evaluations taken directly from the human-robot system. Human-in-the-loop optimization has enabled substantial performance improvements for exoskeleton devices evaluated in a laboratory setting [7], [8], [9], [10], [11] and the real world [12].

Setting up a human-in-the-loop optimization experiment requires the researcher to make many design decisions. Significant time and resources are required to experimentally explore these decisions, which include selecting the optimization algorithm and tuning hyperparameters. Only a few optimization algorithms have been evaluated in existing studies: Covariance Matrix Adaptation Evolutionary Strategy [7], [11], [13], [14], [15], Bayesian optimization [8], [9], [16], [17], [18], gradient descent methods [19], [20], and reinforcement learning [21], [22]. Additional research is needed to better understand the trade-offs between these algorithms and provide methods for selecting the best hyperparameters for a given experimental optimization problem.

Simulating human-in-the-loop optimization provides rapid and direct comparisons between algorithms and hyperparameter settings with fewer experimental resources. Modeling the experimental objective function landscape would allow simulation of human-in-the-loop optimization by providing synthetic evaluations of the objective function. While it is not yet possible to predict how a person will respond to robotic assistance with biomechanics simulations [6], we can fit an empirical surrogate model to experimental data that relates assistance parameters to measurements of the underlying objective landscape, as long as there is a consistent relationship between the parameter space and objective function measurements. This synthetic objective landscape could evaluate any parameter set during a simulated optimization. Although the surrogate model may be useful for informing which algorithm or hyperparameters to use for experimental human-in-the-loop optimization, it does not capture the full complexity of the human-robot interaction and experimental pilot optimizations are still essential to ensure that the optimization is properly initialized.

We modeled the metabolic landscape of people walking with a robotic ankle exoskeleton, simulated human-in-the-loop optimization using this synthetic objective landscape, and compared the performance of different optimization algorithms and hyperparameters. We used experimental data from a prior study [23] where subjects performed experimental human-in-the-loop optimization to find the exoskeleton assistance parameters that minimized the metabolic cost of walking. Surrogate metabolic landscape models were fit to the experimental data to learn a mapping from assistance parameters to the metabolic cost of walking, providing a synthetic metabolic landscape. We explored three simulated human-in-the-loop optimization scenarios optimizing assistance for: expert users with a constant metabolic landscape, novice users adapting to assistance with a time-varying metabolic landscape, expert users using devices with different numbers of assistance parameters. Our simulated human-in-the-loop optimization approach may help inform the selection of optimization algorithms and hyperparameters and reduce the experimental optimization time required to identify optimal assistance parameters (Fig. 1).

**Fig. 1.**
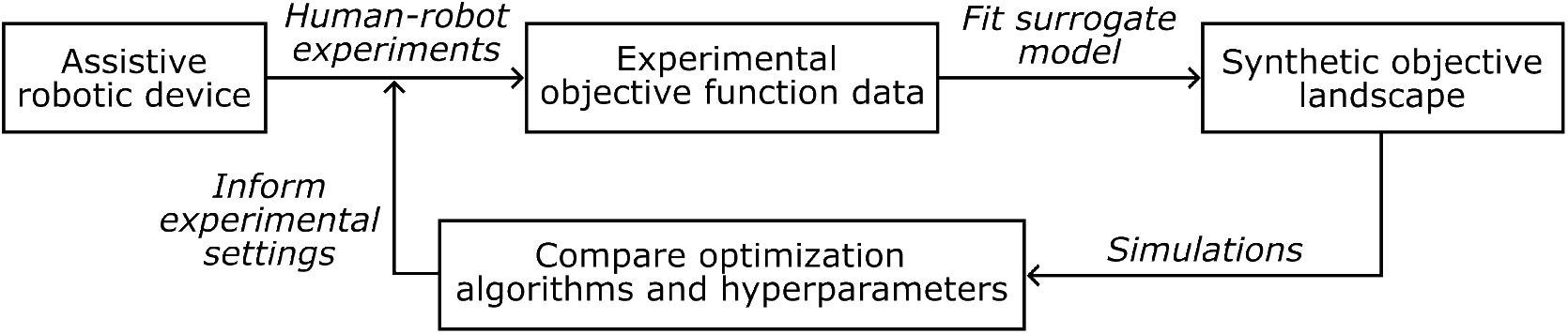
We propose an approach that simulates human-in-the-loop optimization using a surrogate objective landscape model to inform experimental optimization settings. The surrogate is fit from experimental objective function evaluations for different assistance parameter settings, ideally uniformly distributed throughout the parameter space. Evaluations from a range of assistance parameters, ideally uniformly spread over the parameter space, are fit with a surrogate model to provide a synthetic objective landscape. This synthetic landscape enables evaluation of the objective function for any parameter set during simulated human-in-the-loop optimizations. The simulations allow for direct performance comparison between optimization algorithms and hyperparameters to inform experimental human-in-the-loop optimizations.

## II. Methods

### A. Experimental data

Understanding the objective function landscape is important for selecting the appropriate optimization algorithm and hyperparameters to perform human-in-the-loop optimization, but it requires experimental data. We used existing data collected during human-in-the-loop optimization from five subjects walking with an ankle exoskeleton that minimizes the metabolic cost of walking (Fig. 2A)[23]. This previous experiment used the Covariance Matrix Adaptation Evolutionary Strategy algorithm to optimize assistance for six, two-hour sessions with each subject. The dataset contains approximately 180 metabolic cost samples for each subject, a relatively large number of evaluations to cover a space with four dimensions. Assistance parameters were constrained during experimental data collection due to hardware and human safety limitations. Exoskeleton assistance was provided as a torque profile over the gait cycle (Fig. 2B). The torque profile was parameterized with peak torque and peak, rise, and fall times. Peak torque was constrained to 1 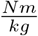 to avoid torques larger than typical biological torques, which could be dangerous for the participant. The peak, rise, and fall times were constrained to be within 35 to 55, 10 to 40, and 5 to 20 percent of the gait cycle, respectively.

**Fig. 2.**
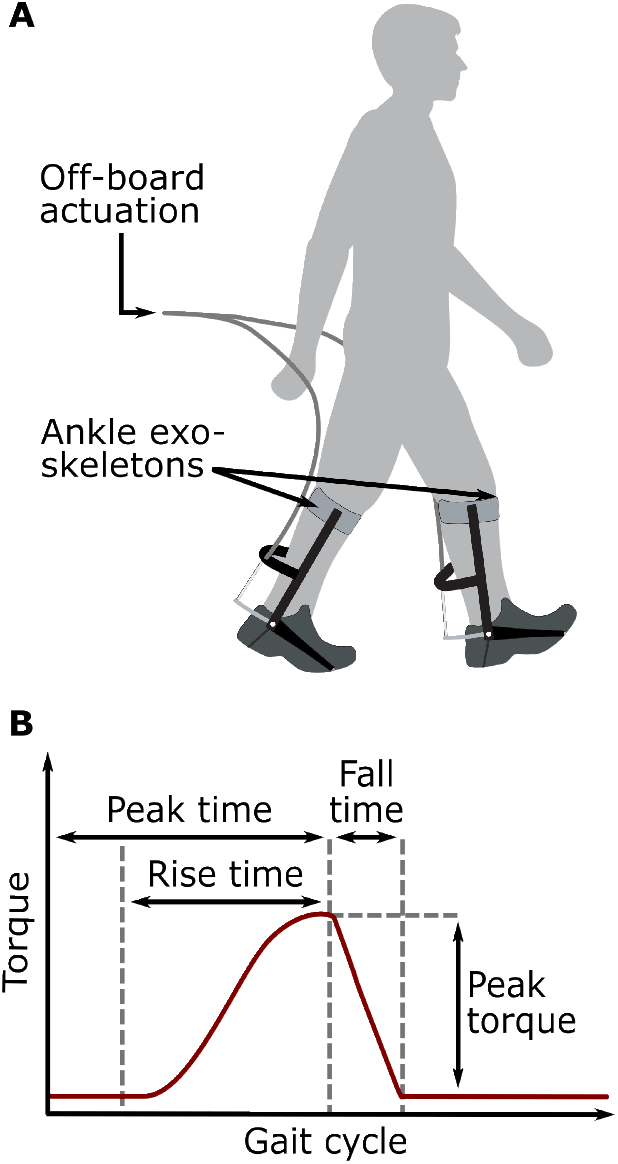
**A**. This study utilizes experimental data collected from participants walking with bilateral ankle exoskeletons [23]. The exoskeletons utilize off-board motors to explore many different assistance profiles without the constraints of portable motors and batteries. **B**. The torque applied at the ankle follows an assistance profile defined by four parameters: peak torque, peak time, rise time, and fall time.

### B. Modeling the metabolic landscape

The metabolic landscape was modeled with linear regression and Gaussian process surrogate models fit to experimental metabolic cost data. Linear regression models used ridge regression with the default regularization parameter of 1. Gaussian process models used constant offset and white noise parameters with a Squared Exponential, Matern, Absolute Exponential, or Rational Quadratic kernel. The parameter and metabolic cost data were normalized across all subjects before being used to train and evaluate the models. The 0.632 bootstrap method was used to evaluate the accuracy of the surrogate models [24]. This bootstrap approach randomly samples the data into training and test sets, trains a model on the training set, and then uses a weighted evaluation with the test set to assess model performance.

### C. Optimization algorithms

We compared several optimization algorithms: Covariance Matrix Adaptation Evolutionary Strategy (CMA-ES), Cross-Entropy, Bayesian Optimization, and Exploitative Bayesian Optimization.

CMA-ES is a generative, model-free, and sample-efficient local optimization algorithm. CMA-ES performs a number of objective function evaluations to collect one generation worth of evaluations. After each generation, CMA-ES fits an *N* -dimensional multivariate Gaussian distribution to the best performing half of the evaluations, the elite samples. Each parameter set, *x*, is drawn from the current Gaussian distribution: 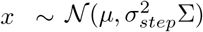, where *μ* is the mean of the distribution that represents the estimate of the parameters with the minimum metabolic cost, *σ*_*step*_ is the step-size scalar that controls the spread of the distribution, and Σ is the covariance matrix [24]. The number of assistance parameters defines the number of dimensions, *N* . The simulated optimizations in this paper use the same optimization hyperparameters as the original experimental data collection and 4 + *floor*(3 ln(*N* )) evaluations per generation [23]. The initial Gaussian distribution was defined with the mean of the parameter boundaries and a step-size scalar value of approximately 30% of the normalized parameter range, the default setting [24].

Cross-Entropy is a common generative algorithm that uses a similar model-free approach to CMA-ES but is less sample-efficient. Each generation of evaluations is used to fit an *N* -dimensional multivariate Gaussian distribution. The number of evaluations per generation and the number of elite evaluations were selected to be twice that of CMA-ES. The initial mean and variance values were set to the same as CMA-ES. These hyperparameters were determined based on a sweep evaluated with one of the metabolic landscape models.

Bayesian optimization is a sample-efficient, global optimization algorithm that balances exploration and exploitation. The objective landscape is modeled with a Gaussian process. All available evaluations are used to fit the prior distribution that captures the believed behavior of the landscape. The prior distribution was initialized with a uniform distribution. The posterior distribution is used to select the next evaluation point to inform sequential samples. We refer to our specific implementation as Bayesian Optimization. We used a Matern kernel to model the Gaussian process and the upper confidence bound as our acquisition function, with default exploration constant of 2.6. The upper confidence bound acquisition function selects the next evaluation such that the upper confidence bound (UCB) is maximized, where *UCB*(*x*) = *μ*(*x*) + *ασ*(*x*). *μ*(*x*) is the mean of the current Gaussian process for the given parameter set, *x. σ* is the standard deviation of the prior distribution at *x* and *α* is the exploration constant [24]. As *α* increases, the upper confidence bound is more likely to explore portions of the parameter space with a higher standard deviation. Lower values of *α* are more likely to exploit the prior distribution knowledge to evaluate the best performing portions of the parameter space with the lowest metabolic cost.

We also compare Exploitative Bayesian Optimization, a Bayesian optimization algorithm that uses the same hyperparameters as Bayesian Optimization except for the exploration constant, which we set to 0.93. This is approximately one-third of the default value and favors exploiting the estimated optimum parameters rather than exploring parts of the parameter space with higher uncertainty. We used a publicly available implementation Bayesian optimization package in Python for both Bayesian algorithms [25].

### D. Simulating human-in-the-loop optimization

We simulated three scenarios of human-in-the-loop optimization, using the lowest-error Gaussian process metabolic landscape model to compare optimization algorithm performance. The scenarios were: 1) optimizing assistance for an expert subject with a constant metabolic landscape, 2) optimizing assistance for a novice subject who is adapting to the exoskeleton and has a time-varying metabolic landscape, and 3) optimizing assistance for an expert subject with parameter spaces of different dimensions. Evaluations were constrained to the same parameter limits used during data collection. Each optimization algorithm used its own process of selecting subsequent evaluations. Measurement noise with a standard deviation of 4.6% was added to every evaluation of the synthetic metabolic landscape. This was done to match experimental measurement noise from estimating the first-order fit using two minutes of respirometry data [7].

*Scenario 1*: We simulated human-in-the-loop optimization for an expert subject walking with an ankle exoskeleton. In this scenario, four parameters (Fig. 2B) were optimized using each of the optimization algorithms. The subject was assumed to have sufficient training to be an expert user who is no longer adapting to the exoskeleton and has a constant metabolic landscape throughout the simulated optimization. A total of 40 simulations were performed for each algorithm, 8 simulations for each of the five subjects. Each simulation consisted of 180 parameter set evaluations.

*Scenario 2*: We simulated human-in-the-loop optimization for a novice subject adapting to the ankle exoskeleton to assess optimization algorithm performance for a time-varying objective landscape. In this scenario, four parameters (Fig. 2B) were optimized using each of the optimization algorithms. Subject adaptation was modeled using a time-varying metabolic landscape. This time-varying landscape linearly interpolated between two different subjects’ metabolic landscape models, starting as the model of one subject and fully interpolating to the second subject’s model after 80 evaluations. The synthetic metabolic landscape was then kept constant for subsequent evaluations. This simulated rate of adaptation is equivalent to 160 minutes walking in the exoskeleton, which is within the range of time it took the subjects from the experimental dataset to reach expert status [23]. A total of 95 simulations were performed for each algorithm, 19 simulations for each of the five subjects. Each simulation consisted of 160 parameter set evaluations.

*Scenario 3*: We simulated human-in-the-loop optimization for an expert subject with 4, 12, or 20 assistance parameters to assess how optimization algorithms perform with larger parameter spaces, which arise for different assistive devices. For example, 6 [18] and 7 parameters [10] have been used for hip exoskeletons, while a hip-knee-ankle exoskeleton used 22 parameters [11]. For our simulations, we selected 12 and 20 parameters to cover this range of existing exoskeleton parameters. To create higher dimensional synthetic metabolic landscapes, we concatenated several of the 4-dimensional constant metabolic landscape models previously derived from experimental data (Fig. 4). We chose to concatenate the data-driven landscape models instead of using a different objective function for the new parameters so they still represent a metabolic landscape that captures an underlying physiological relationship between assistance and metabolic cost. For the 4-parameter case, 8 simulations were performed for each subject for a total of 40 simulations per algorithm. Each simulation consisted of 180 parameter set evaluations. For the 12-parameter case, 10 simulations were performed for each subject for a total of 50 simulations per algorithm. Each simulation consisted of 260 parameter set evaluations. For the 20-parameter case, 5 simulations were performed for each subject for a total of 25 simulations per algorithm. Each simulation consisted of 260 parameter set evaluations.

## III. Results

### A. Modeling the metabolic landscape

The Gaussian process with Absolute Exponential kernel most accurately represented the experimental metabolic landscape and was selected as the metabolic landscape model for our human-in-the-loop optimization simulations (Fig. 3). The contour plots of the Gaussian process models with an Absolute Exponential kernel visually match the five subjects’ experimental data (Fig. 4). Multiple 2D plot projections are used to visualize the metabolic landscape since it is defined by four assistance parameters. The parameter values corresponding to a minimum metabolic cost are different for each subject, which highlights the importance of providing personalized assistance.

**Fig. 3.**
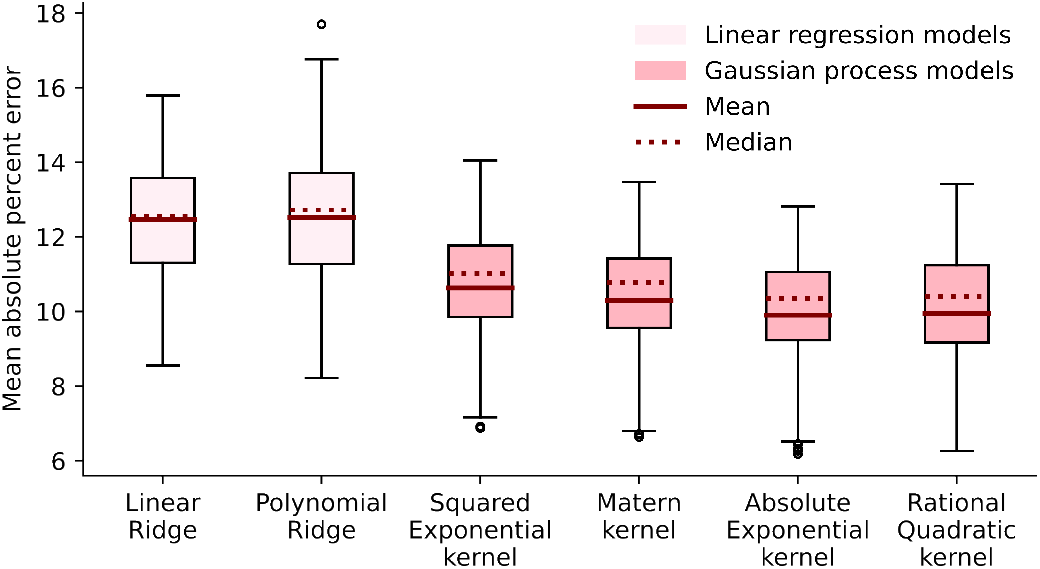
Several probabilistic surrogate models were compared to evaluate how well they fit the metabolic cost data from the ankle exoskeleton experiments.

**Fig. 4.**
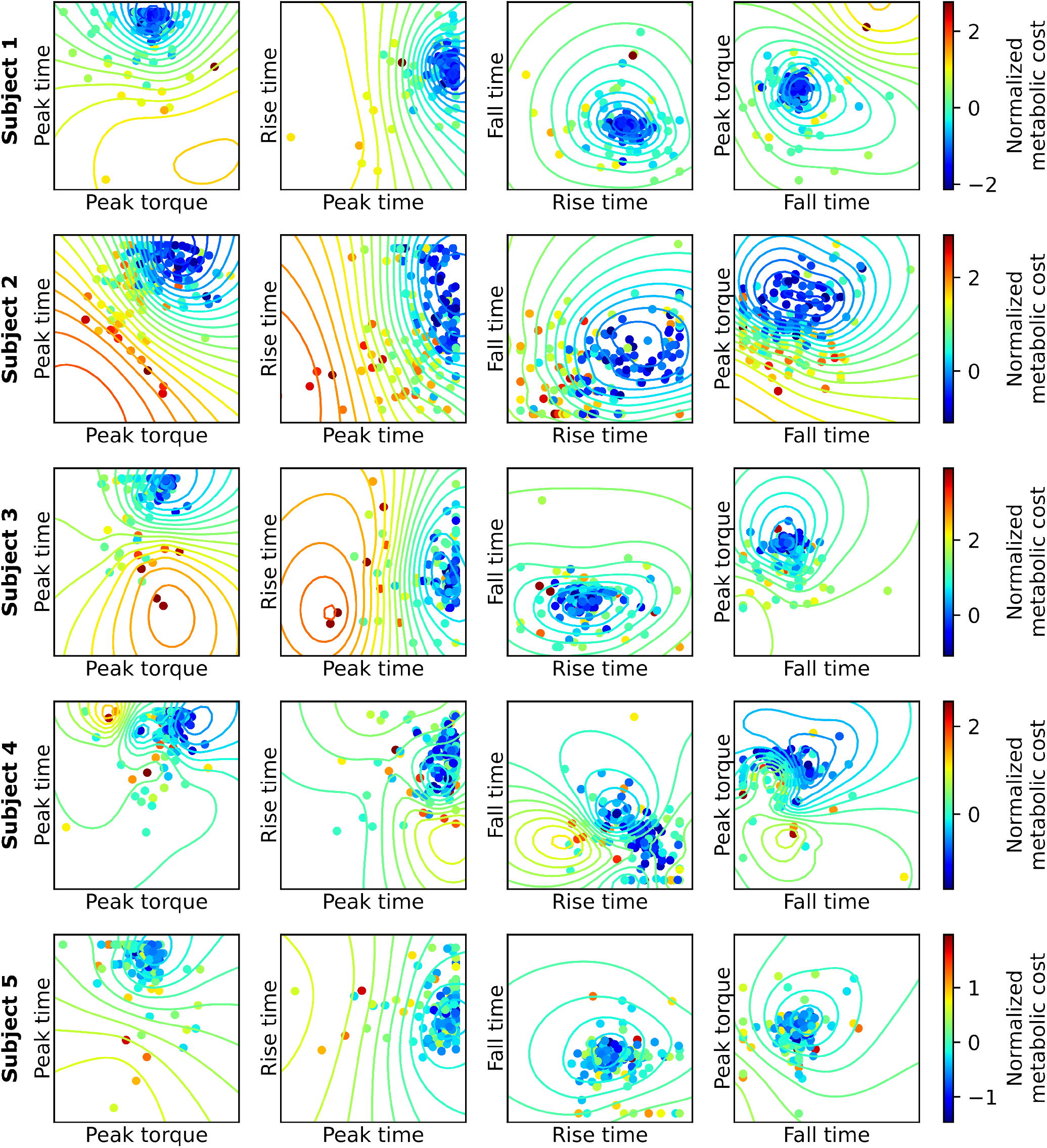
These contour plots represent the metabolic landscapes modeled by a Gaussian process surrogate model with an Absolute Exponential kernel for each of the five subjects. The experimentally-collected data points are overlaid to provide a visual assessment of how well the model captures these data. The data were collected during a human-in-the-loop optimization experiment with the CMA-ES algorithm and are not uniformly distributed throughout the parameter space, causing the surrogate model to be inaccurate in parts of the parameter space with few evaluations. The locations of minimum metabolic cost are different for each subject, highlighting the importance of providing personalized assistance.

### B. Simulating human-in-the-loop optimization

Bayesian optimization algorithms find the optimal assistance parameters most quickly during simulated human-in-the-loop optimizations for expert participants with a constant metabolic landscape (Fig. 5). Bayesian Optimization and Exploitative Bayesian Optimization converge near the optimal solution after approximately 60 evaluations. CMA-ES trends asymptotically to the optimal solution more slowly than the Bayesian algorithms. Cross-Entropy does not reach the optimal parameters and exhibits the largest-variance performance for all simulated optimizations.

**Fig. 5.**
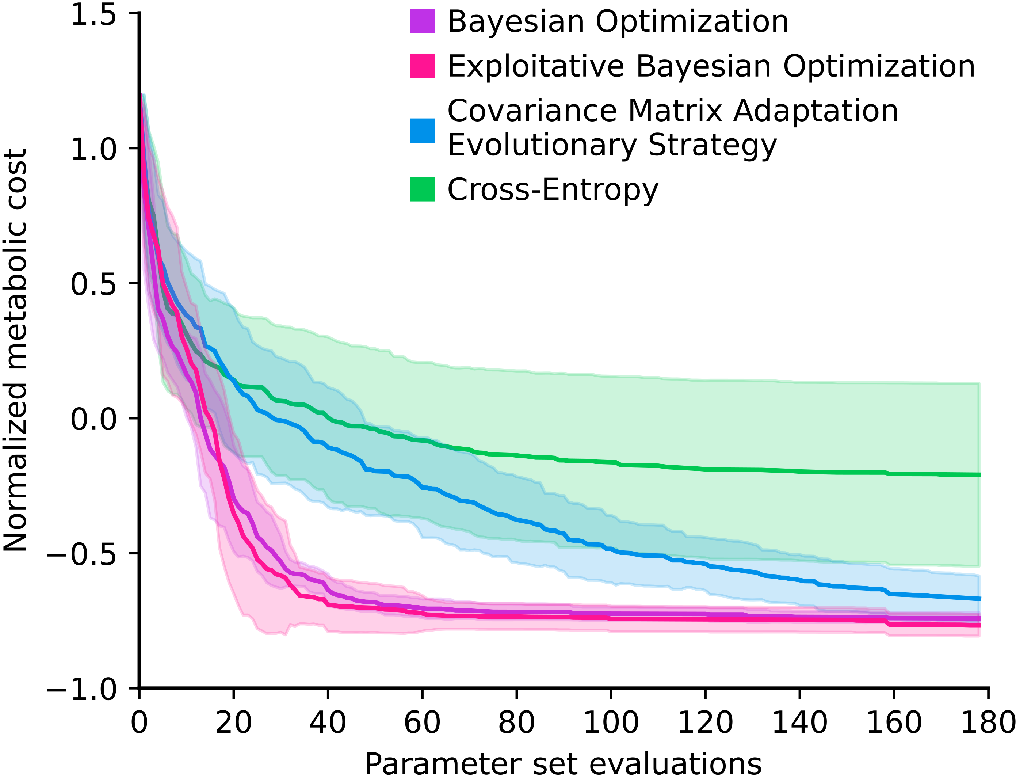
Scenario 1 simulated optimization of assistance parameters for an expert subject with a constant metabolic landscape. Evaluations from the surrogate metabolic landscape included measurement noise with a standard deviation of 4.6% to approximate the experimental collection noise. The line represents the mean value of the minimum metabolic cost up to that evaluation point in the simulation, averaged across all simulations. The error band represents one standard deviation. Each optimization algorithm was tested with 8 simulations for each of the 5 participants. Bayesian optimization methods converged to the optimal parameters with the fewest number of evaluations.

When simulating human-in-the-loop optimization for novice subjects, the time-varying metabolic landscape affects algorithm performance, especially for Bayesian optimization algorithms (Fig. 6). Exploitative Bayesian Optimization finds the optimal solution in approximately 100 evaluations, shortly after the metabolic landscape stops varying at 80 evaluations. Bayesian Optimization trends asymptotically toward the same solution as Exploitative Bayesian Optimization but converges more slowly. CMA-ES converges toward the same optimal solution as the Bayesian algorithms, with similar performance to Bayesian Optimization after 80 evaluations. CMA-ES reaches the optimal solution at a similar rate during both the expert and novice subject simulations. Simulation results for CMA-ES exhibit similar trends to the experimental subject data collected from novice subjects as they adapted to the exoskeleton during human-in-the-loop optimization experiments with a CMA-ES optimizer.

**Fig. 6.**
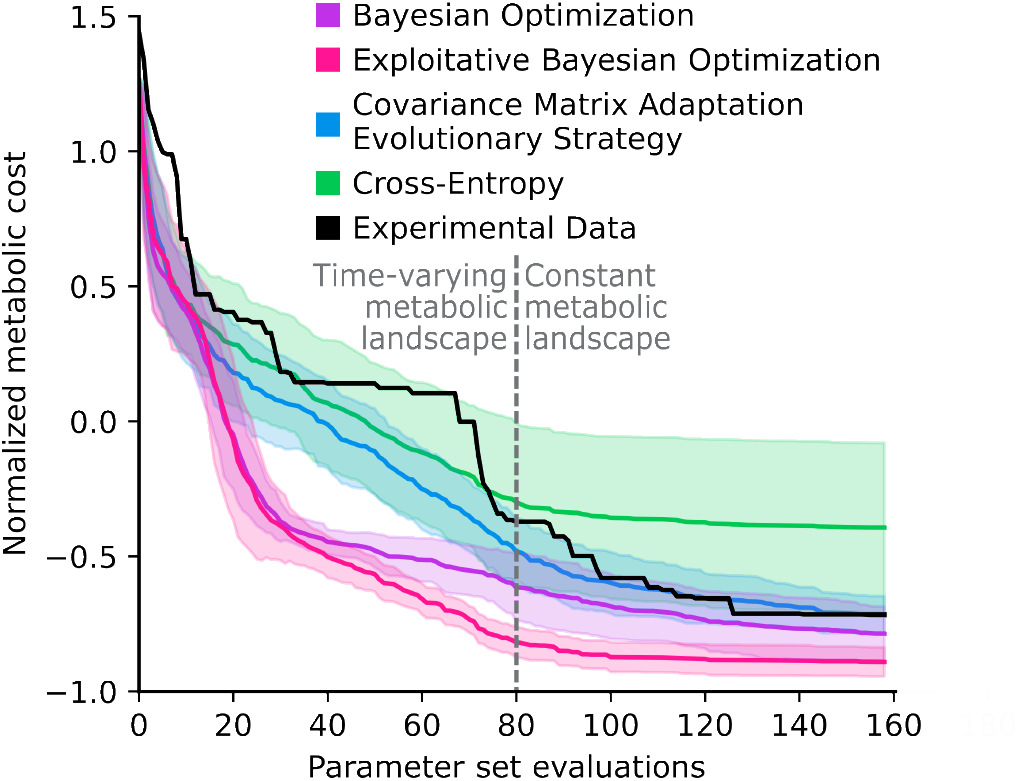
Scenario 2 simulated an optimization of assistance parameters for a novice subject that adapted to the exoskeletons and had a time-varying metabolic landscape. The time-varying metabolic landscape was interpolated between two different subjects’ metabolic landscape models for the first 80 evaluations and then held constant. Evaluations from the surrogate metabolic landscape included measurement noise with a standard deviation of 4.6% to approximate the experimental collection noise. The line represents the mean value of the minimum metabolic cost up to that evaluation point in the simulation, averaged across all simulations. The error band represents one standard deviation. Each optimization algorithm was tested with 19 simulations for each of the 5 participants.

When simulating human-in-the-loop optimization for an expert subject with 4, 12, or 20 parameters, the increased parameter space impacts algorithm performance (Fig. 7). As the number of parameters increases, it is unclear which algorithm is likely to perform the best. Bayesian Optimization reaches the lowest metabolic cost value in the 250 evaluations of the 12- and 20-parameter simulations, but additional simulations and an extended number of evaluations are necessary to further explore this result. Exploitative Bayesian Optimization converges quickly to a suboptimal solution with a larger number of parameters. CMA-ES converges asymptotically toward the optimal solution but does not reach the minimum metabolic cost within the number of evaluations in our simulations. Cross-Entropy converges slowly and does not reach the optimal solution in any simulation setting.

**Fig. 7.**
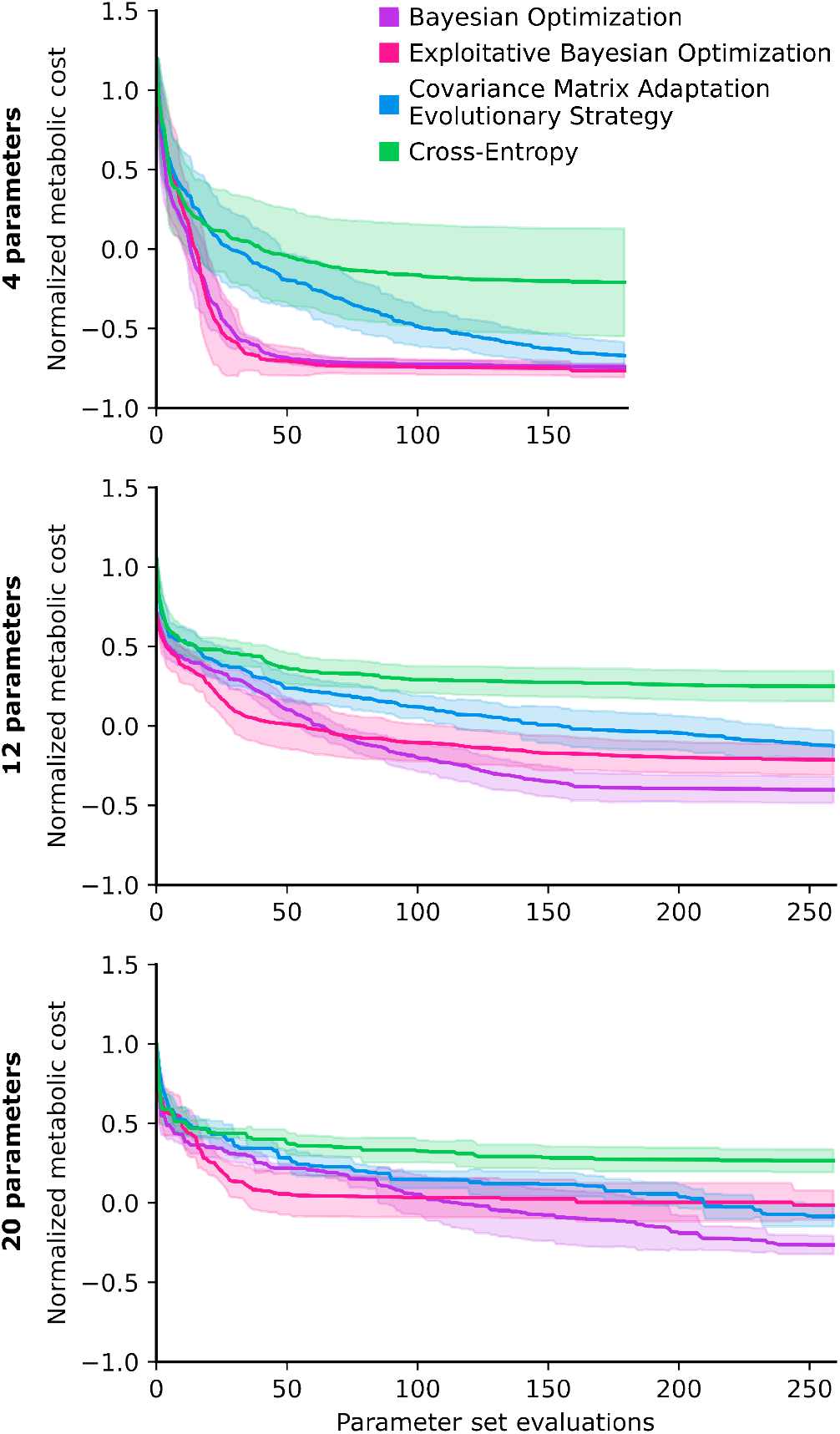
Scenario 3 simulated optimizing 4, 12, and 20 assistance parameters for an expert subject with a constant metabolic landscape. Evaluations included measurement noise with a standard deviation of 4.6% to approximate the experimental collection noise. The line represents the mean value of the minimum metabolic cost up to that evaluation point in the simulation, averaged across all simulations. The error band represents one standard deviation. Each algorithm was tested with 40, 50, and 25 simulations for the spaces with 4, 12, and 20 parameters, respectively.

## IV. Discussion

### A. Modeling the metabolic landscape

Researchers should select the type of surrogate model to fit their objective landscape based on their specific experimental dataset. The Gaussian process models had average errors of 10% on the ankle exoskeleton dataset (Fig. 3), which are relatively low given that the measurement noise for first-order metabolic cost estimates has a standard deviation of approximately 5% [7]. The Gaussian process models are more expressive than linear regression models and better capture non-linearities in the metabolic landscape. However, this increased expressivity requires the experimenter to select additional parameters associated with the kernels. We found many of the Gaussian process model kernels performed similarly and could have effectively captured the metabolic landscape. We suggest comparing several surrogate models on an experimental dataset before identifying the best-performing model.

The experimental data were collected during human-in-the-loop optimization with CMA-ES [23], which performed most evaluations in regions of lower metabolic cost, rather than in a uniform distribution over the parameter space (Fig. 4). The sparse evaluations in regions of higher metabolic cost may reduce the accuracy of the surrogate model and not capture the underlying objective landscape. Ideally, data would be collected uniformly over the parameter space using a grid search. We recommend that researchers perform a grid search for initial data collection to inform objective function models for future simulation studies.

### B. Simulating human-in-the-loop optimization

The qualitative similarity between the simulated and experimental CMA-ES optimization performance for a time-varying metabolic landscape provides evidence that simulating optimization may capture salient information useful for informing experimental optimization (Fig. 6). The experimental data consists of only five subjects who have visible discrete steps in decreasing metabolic cost during optimization, rather than the smoother average profile from 95 simulated subjects. The few number of experimental subjects could explain discrepancies between the simulated and experimental CMA-ES performance. The similar convergence rate of the simulated and experimental CMA-ES optimization indicates that changes in the time-varying metabolic landscape used in simulation may be similar to the changes in the true underlying metabolic landscape as the novice subject adapts to the exoskeleton. However, we do not fully understand how this metabolic landscape changes during adaptation, and poor simulation assumptions may result in ineffective design of a future experimental human-in-the-loop optimization.

Surrogate objective landscape models and simulations may provide a useful approximation for informing experimental human-in-the-loop optimization, but they do not reflect the full complexity of experimental human-robot interaction. It is challenging to predict or model human-robot interaction because of the human’s unique adaptation, volitional control, central nervous system response, and complex dynamic interaction with the robotic system.

CMA-ES converges at a relatively constant rate when optimizing a constant, time-varying, or higher-dimensional synthetic metabolic landscape. The local search structure of CMA-ES may help it perform well during all of these scenarios because it does not keep track of prior evaluations. When the participant has a time-varying metabolic landscape, each generation of evaluations are used to update the posterior distribution and then forgotten, allowing the estimated optimum to move without being constrained by prior information about the underlying landscape. The inherent exploration-exploitation trade-off of CMA-ES seems effective across the different scenarios. CMA-ES converges at a similar rate when optimizing a time-varying metabolic landscape or a landscape with 20 dimensions without hyperparameter tuning. While CMA-ES may take longer to converge than other optimization algorithms, it may be a more reliable choice for many scenarios.

Optimizing time-varying metabolic landscapes may be challenging for Bayesian optimization because it forms a global surrogate model of the objective landscape that must adapt over time. The Bayesian optimizer uses all past evaluations to fit a surrogate model of the objective landscape. As the metabolic landscape changes over time, the Bayesian optimizer still gives the same importance to previous, potentially inaccurate evaluations as new evaluations. Including inaccurate evaluations promotes uncertainty in the Bayesian Optimization’s surrogate metabolic landscape model. For example, the surrogate model would have a higher value of uncertainty in a region where there are two evaluations for the same set of parameters with two different objective measurement values. This uncertainty in the surrogate model will alter which evaluation points are selected next based on the upper confidence bound criteria. This uncertainty may cause the optimizer to converge on a suboptimal solution or require more evaluations to reach the optimal solution (Fig. 6).

In the scenario optimizing assistance for a novice user with a time-varying metabolic landscape, Bayesian Optimization required more evaluations to converge to the optimal solution than Exploitative Bayesian Optimization. Bayesian Optimization has a higher exploration constant and may have performed evaluations spread across the parameter space, providing less frequent evaluations near the time-varying optimum and requiring more evaluations to reach the optimal solution. Exploitative Bayesian Optimization attempts to overcome this challenge by having a lower exploration constant that favors exploiting its surrogate model by more frequently evaluating near the best performing portions of the parameter space rather than exploring more uncertain parts of the space. Exploitative Bayesian Optimization may converge to the optimal solution with fewer evaluations because the more frequent evaluations near the estimated optimum parameters closely tracks changes as the optimum varies with time. Selecting an appropriate exploration constant value may be important for successfully optimizing time-varying landscapes with Bayesian optimization. Simulating human-in-the-loop optimization may help inform what exploration constant value may perform well in experiments. Although, experimental piloting is important to capture the full complexity of human-in-the-loop optimization and validate any simulated hyperparameter selections.

It is unclear which optimization algorithm performs the best in higher dimensional parameter spaces. All algorithms converge toward an asymptote more slowly as the number of parameter increases. Bayesian Optimization finds the minimum metabolic cost solution within the evaluations tested in our simulations, but CMA-ES is still improving its best set of parameters at the end of the evaluations. Additional simulations with an increased number of evaluations are necessary to better understand this trade-off between convergence rate and achieving the lowest possible metabolic cost. The performance gap between Bayesian Optimization and CMA-ES decreases as the number of parameters increases. We hypothesize that CMA-ES will perform better than Bayesian Optimization when optimizing more than 20 parameters, especially if other factors like a time-varying metabolic landscape are introduced. Exploitative Bayesian Optimization converges to a suboptimal value in the higher-dimensional parameter spaces, possibly because its lower exploration constant may cause it to keep evaluating parameter sets near a local minimum point it finds early in the optimization.

Results from our three simulated scenarios suggest there is no clearly superior optimization algorithm for human-in-the-loop optimization for all use cases. Selecting the algorithm and hyperparameters based on the unique criteria for each experiment increases the likelihood of performing a successful human-in-the-loop optimization. Experimental designers should consider factors like measurement noise, adaptation of the participant, and number of assistance parameters. In practice, many of these factors are likely to be present in an experiment, making for a challenging optimization problem. We suggest that researchers perform simulated optimizations to inform algorithm and hyperparameter selection and then experimental pilot optimizations to confirm the simulation selections. Simulated optimizations should use relevant experimental data to match the optimization challenges for each specific application as closely as possible. Copying the algorithm and hyperparameters used in a different human-in-the-loop optimization experiment may result in an ineffective optimization.

We provide open-source code to enable others to replicate and extend our results to explore additional optimization algorithms, perform hyperparameter tuning, and extend simulated optimization to datasets for new applications. [26]. Gradient descent is not explored in this project but has been used for human-in-the-loop optimization of a single parameter [19]. Reinforcement learning may be another viable alternative outside of traditional optimization algorithms [21], [22].

## V. Conclusion

Our simulation approach for human-in-the-loop-optimization (Fig. 1) may provide a method to compare different optimization algorithms and hyperparameters for new assistive device applications to accelerate experimental human-in-the-loop optimization. Researchers with a new assistive device could perform initial human-subject experiments to evaluate the objective landscape over the assistance parameter space, ideally with a uniform method like a grid search. Surrogate models could be fit to these experimental evaluations and compared to select the most accurate synthetic landscape model. The synthetic objective landscape obtained from the surrogate model would enable simulation of human-in-the-loop optimization by evaluating parameter sets without experimental measurements. Simulated optimizations could help researchers identify which algorithms and hyperparameters may perform well for their specific application, before conducting experimental human-in-the-loop optimizations. Well-designed simulated optimizations may save researchers experimental time and resources when performing human-in-the-loop optimization experiments for new devices or applications.

